# Vagal nerve stimulation alters task-evoked pupillary responses in older adults but not younger adults in a single-blind sham-controlled crossover trial

**DOI:** 10.1101/2025.10.10.681160

**Authors:** Elizabeth Riley, Genevieve Wager, Samantha Rahman, Eve De Rosa, Adam Anderson

## Abstract

**Introduction:** The locus coeruleus (LC) undergoes age-related changes and is involved in Alzheimer’s disease pathogenesis. Vagal nerve stimulation (VNS) may modulate LC activity and could be used therapeutically, but age-related differences in VNS responses remain unexplored.

**Methods:** We used a single-blind, sham-controlled, crossover design in 41 participants (21 younger, 20 older adults). Participants completed a visual oddball task with pupillometry during transcutaneous auricular VNS (verum: cymba concha; sham: earlobe) with ∼30-minute washout between conditions.

**Results:** Older adults showed smaller baseline pupil diameter but larger normalized task-evoked responses than younger adults a priori. VNS produced age-specific effects: older adults demonstrated increased tonic pupil size throughout stimulation and reduced oddball-evoked responses, with stronger effects with more current. Younger adults showed no consistent VNS effects.

**Discussion:** VNS affects LC-related physiological measures differently across age groups, with older adults showing more robust responses. These age-specific effects may reflect different baseline LC activity states.

## 1 Introduction

The locus coeruleus (LC) of the brainstem plays an important role in Alzheimer’s disease (1–6), Parkinson’s disease (7–10) and frontotemporal dementia (11,12), and potentially other neurodegenerative diseases (13,14). It accumulates pretangle hyperphosphorylated tau starting in the 2nd decade of life (6), and its cells die early on in the progression of several neurodegenerative diseases. In addition to being one of the earliest cell populations vulnerable to AD-related pathology (15,16), it may also spread pathology to subcortical and cortical areas (17,18). It is thought to be one of the primary places in which AD, PD, and frontotemporal dementia begin. Therefore, it is of great interest to assess and preserve its health.

Recent research increasingly underlines the LC’s critical roles in sleep, stress, memory, and attention. A common theme across all of this work is that in order to achieve optimal outcomes in cognition, the LC needs to have an optimal amount of activity - neither too little, nor too much (19–21). Overactivity leads to stress (20,22,23), and underactivity leads to hypoengagement (17,24). Both are associated with aging; nonhuman animal research demonstrates that mild damage to the LC results in hyperactivity (25,26), while human research suggests severe LC damage results in hypoactivity (3). The hope is to uncover ways to modulate locus coeruleus activity that may improve its functioning, reduce its vulnerability, or even potentially limit the propagation of disease.

Research into how to modulate LC activity safely and effectively is consequently a growing area of research. Strategies in humans include the use of various pharmacological agents specific to the LC (27), exercise (28), deep breathing (29), and vagus nerve stimulation (VNS) (30–32). Amongst these methods, VNS is uniquely accessible, requiring only a portable stimulator, and can be used by anybody regardless of age or disability. VNS can be delivered either invasively (with surgical implantation of a stimulator in the chest) or noninvasively, using transcutaneous stimulation at sites on the neck or cymba concha of the ear. However, the ideal stimulation frequency, level of current, and timing of stimulation for LC modulation remain unknown.

The vagal nerve, cranial nerve X, serves as the major pathway by which internal states are communicated with and regulated by the brain (33). The vagus is a critical bridge between the body and the brain, and substantial evidence shows that fluctuating cardiac phases and baroreceptor activity are continuously modulating cognitive and affective function (34,35). As the locus coeruleus is the “second stop” in the afferent vagal pathway, receiving input from the nucleus tractus solitarius which is directly innervated by the vagus, there is a clear anatomical pathway by which breath control, cardiovascular exercise and direct afferent vagal stimulation can modulate LC activity. Numerous reports have already demonstrated the ability of transcutaneous VNS to modulate LC function in humans, although significant heterogeneity in the direction and nature of the effect demands further investigation (36–40). Additionally, the efficacy of VNS for symptoms of cognitive decline in older adults (41–43) lends support to the idea that LC modulation can have a positive impact on the course of neurodegenerative disease. However, there is currently no information about the differences or similarities in the effects of VNS on younger vs. older individuals. Since the LC is known to experience major changes in structure and function with age (44–47), in accordance with its role in the long preclinical phase of AD, it is quite possible that younger and older adults will respond differently to VNS.

Pupil size serves as an external reflection of the effects of VNS on the LC (48). Pupil size is regulated by affective and cognitive states and is a reasonable proxy for human LC activity under the right conditions. Here, we examined pupillary responses in the context of behavioral orienting in a classic visual oddball task, in which context pupil size is especially well linked to LC activity (48). This trial (49) employed a single-blind, sham-controlled, crossover design using transcutaneous auricular VNS. Each participant received both verum and sham stimulation in a single visit, with a ∼30 minute washout between conditions.

## 2 Methods

### 2.1 Participants

We recruited 46 participants, with 24 under age 40, and 22 over age 60 for clinical trial https://clinicaltrials.gov/study/NCT06880510 (49) as shown in Fig. 1. One older adult declined to participate, and one younger adult was not eligible (heart murmur). Data from one younger adult and one older adult had to be excluded because of technical problems with the VNS device. One further older adult had to be excluded because of severe eczema which made VNS painful (this was the study’s only adverse event; the participant’s pain resolved immediately upon cessation of stimulation, which lasted only a few seconds), resulting in a final group size of 21 for younger adults (67% women, mean age = 24.94 years, SD = 6.33) and 20 for older adults (70% women, mean age 71.60 years, SD = 4). Participants completed this trial as part of a larger study that also involved MRI, and thus were screened with exclusion criteria that included MRI safety, neurological disease and cardiac arrhythmias. Testing took place at Cornell University from July 2024 through August 2025, and was covered under IRB protocol 1910009087.

**Figure 1.**
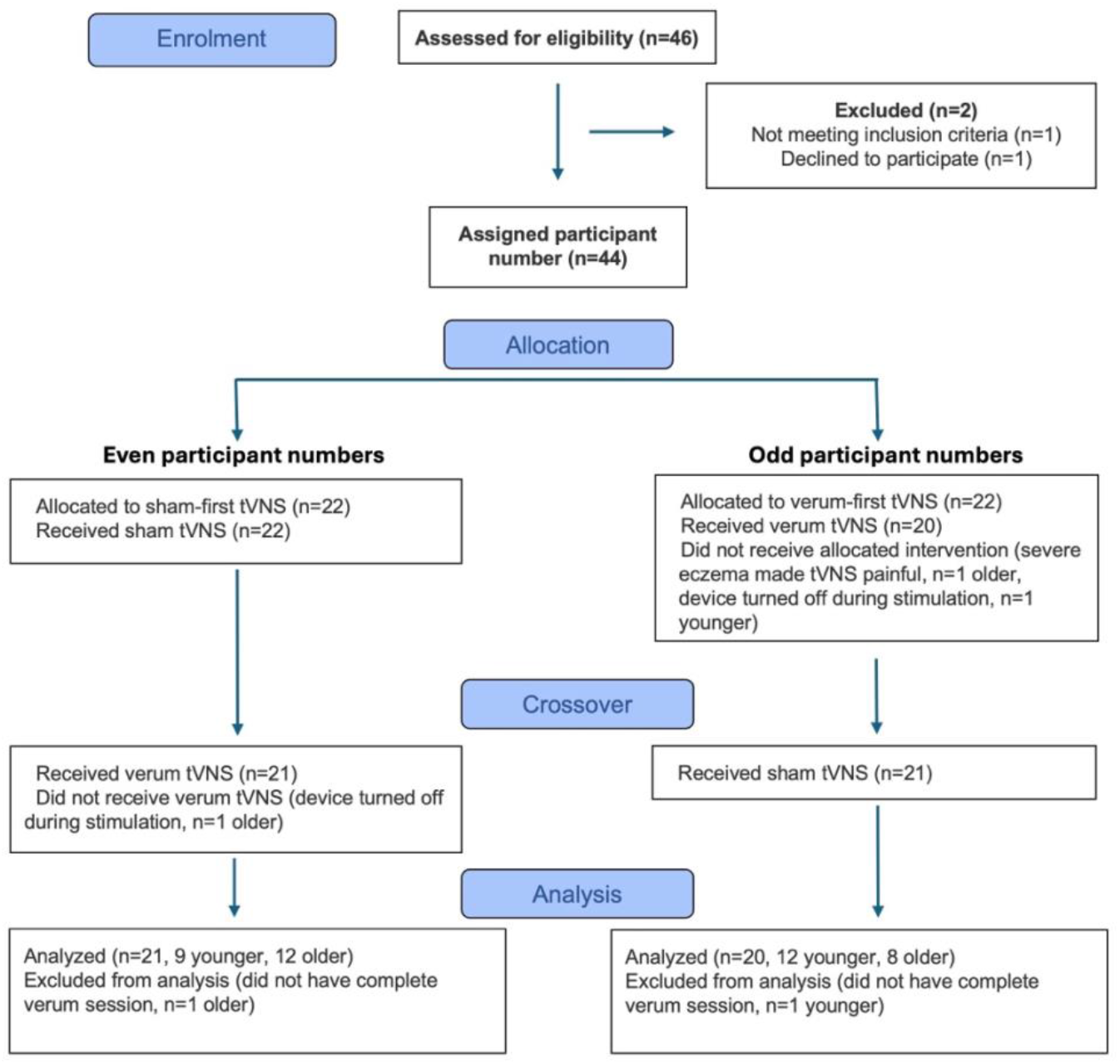
Flow diagram of study enrollment, allocation, crossover and analysis.

### 2.2 Study procedures

First, experimenters obtained informed consent. Each participant completed two sessions of pupillometry during an oddball task with transcutaneous electrical stimulation. In the sham session, the stimulator was placed on the left earlobe, while in the verum session, it was placed at the cymba concha of the left ear. The order of the sham and verum sessions was counterbalanced by the assigned participant number (odd = verum first). The participants were blind to which position was expected to stimulate the vagus nerve. Before each session, the electrical stimulator was attached, and the current level was adjusted individually for strong sensation. The lights were turned off during calibration, which took approximately 3 minutes. Lights remained off during both sessions of the task. The room had no windows. The task took approximately 8 minutes to complete. During the washout period (approx. 30 min), participants undertook cognitive testing and filled out surveys related to other aspects of the overall visit. All data was stored on a secure server managed by Cornell University.

#### 2.2.1 Device and device usage

This trial used a tVNS Technologies research stimulation device, with a Lenovo tablet M8 using the tVNS Technologies tVNS Research app to control settings. Stimulation was set to a frequency of 25 Hz, 250 μs pulse width, and 50% duty cycle (30 seconds on, 30 seconds off). These settings were selected to match previous tests that had shown efficacy previously (50). The left ear was chosen given the laterality of afferent vagal fibers (51), thus reducing efferent effects on the heart. Before placing the device on the ear, participants wiped their outer ear with an alcohol wipe. The stimulator tip was covered in tVNS Technologies Electrode Cream liberally before every application of the device (i.e., twice per participant), and new tVNS Technologies Electrode Pads were applied for each participant.

The stimulation current was adjusted according to each subject’s sensation. The current intensity at the beginning of the adjustment period was 100 µA and was increased by the experimenter in steps of 100 µA until the subject reported a dominant vibrating sensation in the region acute to the electrode. Most commonly, participants reported a pin-prick sensation at lower amplitudes. As the voltage was increased, this generally became more of an itch-like sensation before ultimately becoming a comfortable vibration. Occasionally, increases in voltage would make the sensation less comfortable. To maximize participant comfort and ensure that the stimulation would be tolerable for at least 8 minutes, voltage was again decreased in 100 µA increments until an optimal level was reached in which the stimulation was detectable but not bothersome or painful. Since sensitivity to the stimulation varies from person to person, it is not surprising that some participants never reported feeling a vibrating sensation, or even any stimulation at all. In these cases, we continued to increase the stimulation in 100 µA increments, frequently checking in with participants and ensuring that there were no changes in their experience. Once we reached 5000 µA, the maximum current possible on our stimulation device, we confirmed that there was no discomfort or pain and proceeded to the oddball task.

#### 2.2.2 Pupil and eye movement assessment

Eye movements, blinks, and pupil size were recorded with an Eyelink 1000 Plus eye tracker focused on the right eye at 1000 Hz sampling rate. The eyetracker was fixed in place and was not moved between participants during the entire course of the trial. Participants used a chinrest (SR-Research), which was screwed to the desk, to remain still (eyes 72 cm from the center of the screen). The eye tracker was calibrated just prior to task onset for each person, both times, using a 9-point calibration routine. During both task runs, participants were reminded to blink only when necessary for comfort and center their gaze on a fixation dot.

Raw pupil data for each participant was formatted using the EyeLink DataViewer application (SR-Research), converted to mm using calibration factors calculated using measurements from an artificial eye as per SR-Research instructions, and then cleaned using custom routines in MATLAB and R. Any blinks flagged by EyeLink software, plus inside a 25 ms margin before and after the blink, were removed. Since blink artifacts were still present after filtering those automatically flagged, remaining blink artifacts were then removed, with margin (and subsequently interpolated), by thresholding the data using a formula recommended by Kret and Sjak-Shie (52): (median normalized dilation speed) + 8 * (median absolute deviation). The factor of 8 was chosen empirically to fit our needs as recommended by Kret and Sjak-Shie. Data was then interpolated to replace blinks using the *approx* function in R.

After preprocessing, participants’ pupil size dynamic range was estimated by calculating the difference between the 1st and 99th percentile clean pupil diameter values for that individual across the entire sham run. For analysis by trial, pupil size was normalized by subtracting the average pupil size (mm) of the 50 ms before the onset of the trial from all values in the trial (i.e., baseline normalization), and then divided by the dynamic range. The pretrial pupil size was also used to index baseline pupil diameter, an putative index of tonic LC activity.

To assess the effect of VNS condition on tonic pupil size, we first calculated the trialwise difference between pretrial baseline pupil size under sham and verum conditions for each participant. This was necessary because of drift in baseline pupil size throughout the course of the experiment, likely reflecting decreased arousal. In addition to calculating an average pupil size difference between conditions for each participant, we also calculated the average difference at each of 6 equal epochs (25 trials, ∼ 1.3 min each) throughout the task.

#### 2.2.3 Task details

The oddball task had 3 types of stimuli: standard (60%, 90 trials), oddball (20%, 30 trials), and bright (20%, 30 trials). Standard stimuli were 1.8° degrees visual angle purple circles, while oddball stimuli were 3.6° visual angle purple circles, both presented on a gray isoluminant background. Oddball stimuli required a button press on a standard keyboard. Bright stimuli were the same size and color as standard stimuli, but were brighter than standard stimuli. Participants were not told about the bright stimuli. All stimuli were presented for 75 ms each. Between stimuli, participants fixated on a black fixation dot (0.15° visual angle). The inter-stimulus interval was jittered between 2.5-3.5 s (uniform). The order of trials was pseudorandom such that neither oddball nor bright stimuli occurred within two trials of each other. Total task time was approximately 8 minutes. The task was modeled on the task described by Murphy et al. (48) to ensure maximum relevance to LC functioning.

### 2.3 Statistics

All models were constructed in R using the *lme4* package (53). For linear mixed effects models, *lmer* was used, except for analyses in which the outcome variable was a single average per participant (e.g. overall task performoance), in which case a random effect of participant ID was not needed and *lm* was used for simple linear models. Models were evaluated using *joint_tests* from the *emmeans* package. All post-hoc tests were done using *emmeans*, with p values adjusted for multiple comparisons using the Tukey method. Partial eta squared effect sizes were calculated using the *emmeans* package also, while Cohen’s d was calculated directly from marginal means and the model pooled standard deviation.

### 2.4 Power analysis

We considered the effect size of transcutaneous VNS in the context of an oddball task administered in a crossover design, the closest study design to this one available. In this study (54), an η^2^_p_ = 0.13 was reported in a sample size of 20. To detect an effect of this size at 90% power with alpha = 0.05 in a paired sample (crossover design), a sample size of 19.6 is required according to the *pwr* package in R. We therefore sample size of 20 per group, or 40 total.

### 2.4 Open science policies

De-identified participant data and analysis scripts will be available on OSF (osf.io/vz85h/). All results are distributed directly to participants through a study newsletter with plain language summaries, and through accessible lectures for the public.

## 3 Results

### 3.1 Task performance

Participants performed well on the oddball task, with younger adults correctly identifying 99.07% (SD = 1.17) of oddballs, and older adults identifying 99.33% (SD = 0.99). Young made errors of commission 0.97% (SD = 1.39) of the time for standard trials and 0.20% (SD = 0.59) of the time for bright trials, and older adults made such errors 0.43% (SD = 0.89) and 0.19% (SD = 0.84) of the time respectively. There was no significant difference in task performance between groups (t = -0.008, p = 0.993). Task reaction time was significantly faster in younger adults (425.89 ms, SD = 83.23) compared with older adults (472.55 ms, SD = 113.41, t = 7.81, p < 0.0001).

### 3.2 Baseline pupil diameter

Pretrial baseline pupil diameter, our measure of tonic pupil size, was larger in younger adults (3.68 mm, SD = 0.72) than in older adults (3.35 mm, SD = 0.53, t = -19.30, p < 0.0001, η_p_^2^ = 0.06, Fig. 2a), consistent with age related myosis. Dynamic range of pupil size was also larger in younger adults (1.50 mm, SD = 0.71) than older adults (0.90 mm, SE = 0.27, F(1,226) = 75.86, p < 0.0001, η_p_^2^ = 0.25), replicating prior findings (44). Pupil diameter decreased throughout each 8-minute task run (150 trials per run) in both age groups (Fig 2b).

**Figure 2.**
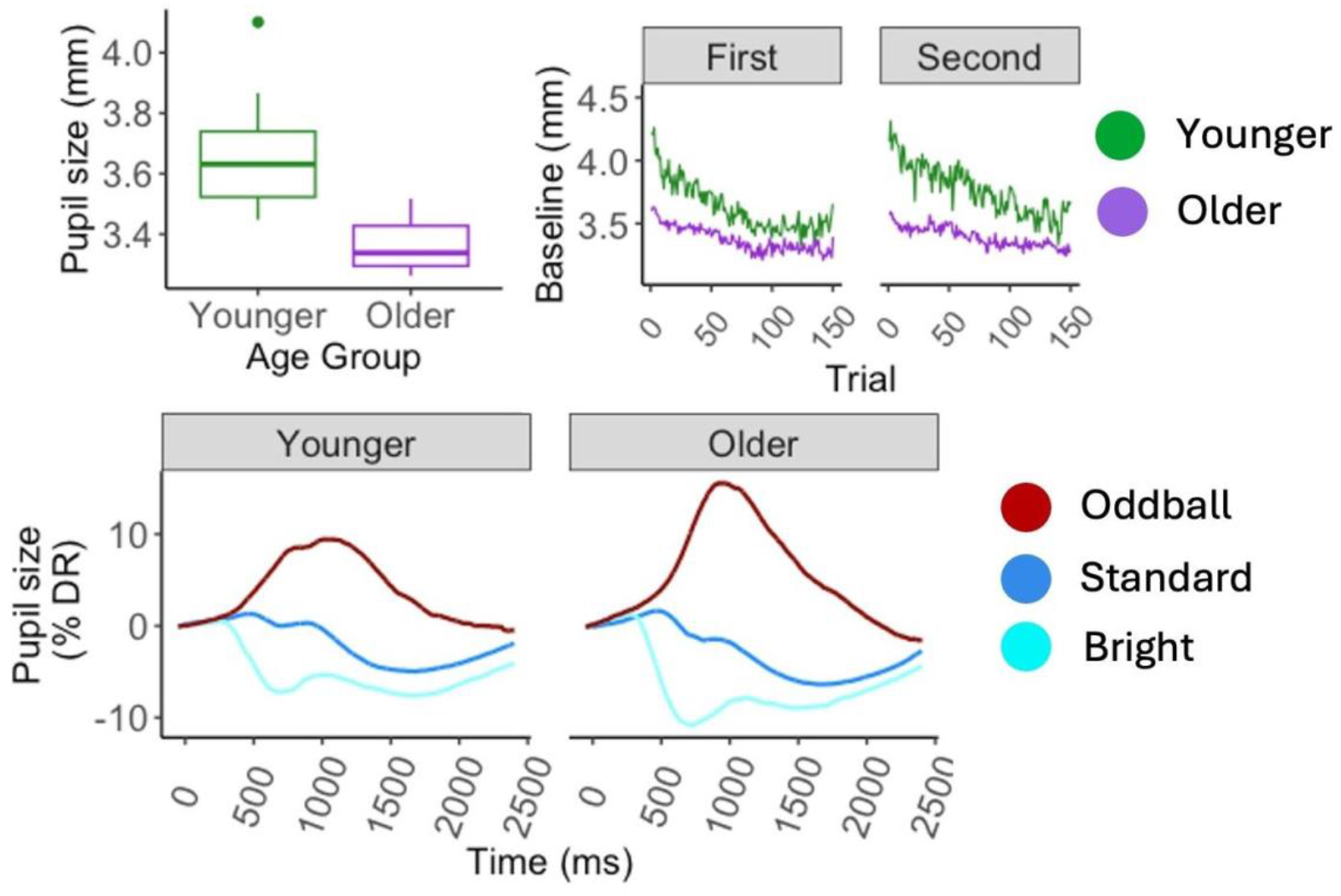
(a) Boxplot of individual average pupil sizes in mm under sham conditions by age group. Box boundaries represent 25th and 75th percentiles, center line shows median, and whiskers extend to 1.5× IQR or the most extreme data point within that range. Points beyond whiskers are outliers. (b) Average pupil size in mm by trial, age group and session. The 150 trials were completed in ∼8 min. (c) Task-evoked pupillary responses expressed as percent of individual dynamic range, by age group and trial type. Oddball trials required a button push. Standard error is too small to be visible.

### 3.3 Phasic pupillary responses

To assess the effect of oddball task conditions on phasic pupillary responses, we fit linear mixed-effects models on pupillary response maxima (minima, for bright trials) under sham VNS conditions. Fixed effects included age group and trial type, along with their interaction. Participant ID was included as a random intercept. The model was evaluated using joint tests. There was a significant effect of trial type (F(2,Inf) = 508.65, p < 0.0001, η_p_^2^ = 0.49), a marginal effect of age group (F(1,Inf) = 3.36, p = 0.067, η_p_^2^ < 0.001, and a significant interaction between age and trial type (F(2,Inf) = 28.92, p < 0.0001, η_p_^2^ = 0.008). Pairwise comparison of marginal means showed that each of the three trial types differed significantly from the others (all p < 0.0001) within both age groups. Oddball trials resulted in robust pupil dilation relative to standard trials. Bright stimuli resulted in clear pupil constriction relative to standard trials, consistent with the pupillary light reflex. Older adults had significantly larger oddball responses than younger adults in units of percent dynamic range (z ratio = -3.85, p = 0.0016, d = 0.29), as well as significantly larger pupillary light reflex pupil constrictions (z ratio = 3.89, p = 0.0014, d = 0.29), as illustrated in Fig. 2c. Pupil constriction minima to bright trials occurred more rapidly than pupil dilation maxima to oddball trials, likely reflecting with the more cognitive nature of behavioral orienting to stimulus salience (Fig. 2c) with similar temporal dynamics in younger and older participants.

We also repeated the linear mixed effects model on pupillary response maxima including a fixed effect of pretrial baseline. The results of the model were the same as for the previous model, with the addition of a significant effect of pretrial baseline such that larger baseline predicted smaller phasic maxima (F(1,Inf) = 1051.912, p < 0.0001, η_p_^2^ =0.11). Because an inverted-U or quadratic relationship between pretrial baseline and phasic maxima has been reported, we repeated this model with a quadratic term for baseline, and compared it with the model with a linear term for baseline. The model with a quadratic term was a worse fit (log likelihood with linear term 4678, log likelihood with quadratic term, 4583). We therefore used a linear term for subsequent models.

### 3.4 VNS administration parameters

Transcutaneous auricular VNS was administered to each participant at sham (earlobe) and verum (cymba concha) positions, in a single-blind manner with order counterbalanced, with the current individually adjusted to strong sensation as per 2.2.1. The average current used during sham stimulation was 1978.18 μA (SD = 1121.95), and the average current used during verum stimulation was 2073.46 μA (SD = 857.58). Current did not differ significantly by age group or VNS condition (for age group, t = 1.64, p = 0.11, for VNS condition, t = 0.52, p = 0.60). The average amount of time between administrations was 27.90 min (SD = 7.53) for younger adults and 37.89 min (SD = 6.52) for older adults, which was a significant difference (t = 6.14, p < 0.0001). This is because in between VNS conditions, participants were given cognitive tests, which took longer to administer in older adults than younger adults.

#### 3.5 Effect of VNS on task reaction time

To assess the effect of verum vs. sham VNS (i.e., VNS condition) on task reaction time, we fit a linear mixed-effects model on reaction time with fixed effects of VNS condition, condition order and age group and their interaction, and with a random slope for VNS condition and a random intercept for participant ID to account for the crossover design. The model was evaluated using joint tests. There was again again a significant effect of age group (F(1,34) = 8.27, p = 0.0069, η_p_^2^ = 0.20), but no significant effect of VNS condition (F(1,33.98) =0.032, p = 0.86), or order (F(1,34), = 0.044, p = 0.84), nor any significant interactions.

### 3.6 Effect of VNS on tonic pupil size

In a simple linear model, with average tonic pupil size difference (Fig. 3) per participant across all trials as the response variable, and age group and condition order as fixed effects, we found a significant effect of order (F(1,34) = 6.79, p = 0.013, η_p_^2^ = 0.16), and a significant interaction between age group and order (F(1,34) = 5.16, p = 0.029, η_p_^2^ = 0.13). Examining marginal means, we found that older adults had an estimated positive pupil size difference (i.e., pupil size increased with verum VNS) in both condition order groups and that these did not differ from each other (t ratio = -0.62, p = 0.93). However, younger adults had a significant order effect (t ratio = -8.714, p < 0.0001, d = -1.67), entirely consistent with an effect of session, rather than an effect of VNS condition.

**Figure 3.**
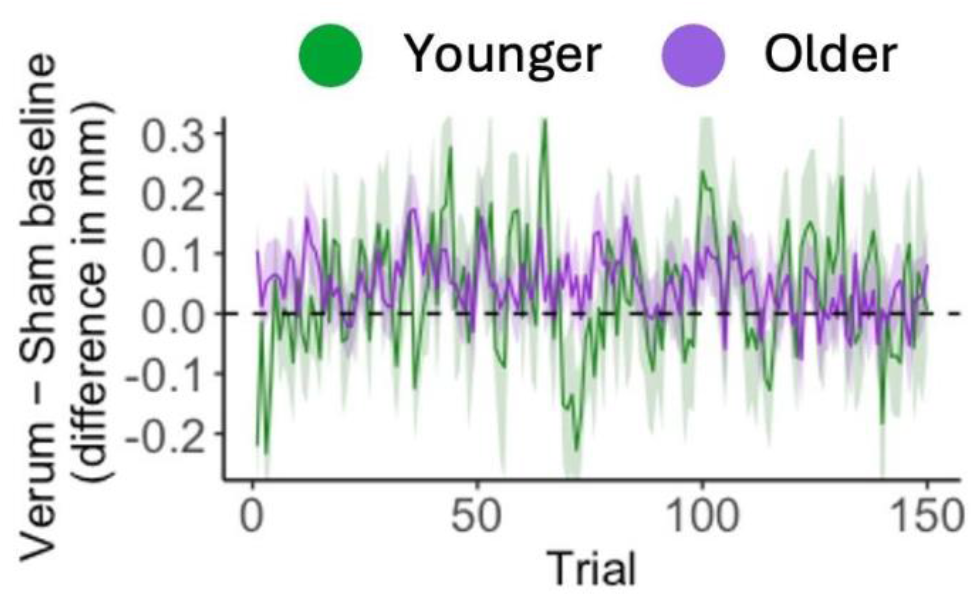
Difference in mm between verum and sham pretrial baseline pupil size at each trial throughout the course of the oddball task, by age group. Ribbon shows standard error of the mean. Positive values reflect an increase in pupil size with verum stimulation.

To investigate how the effect of VNS on tonic pupil size might change throughout the course of the 8-minute stimulation, we tested for differences in pupil size within each of 6 75s epochs (as described in 2.2.2). We conducted one-sample t-tests within younger and older adult groups separately at each of the 6 timepoints, and then adjusted all p values for multiple comparisons using the FDR method. In younger adults, there was no consistent pattern. In older adults, tonic pupil size difference was significantly increased during all epochs (all p < 0.001), except for the final one and thus 83% of the stimulation session.

### 3.7 Effect of VNS on dynamic range

We tested whether VNS affected dynamic range using a linear mixed effects model with fixed effects of age group, VNS condition, and condition order and their interactions, and a random intercept for participant ID. We found the expected effect of age group (F(1,68) = 31.61, p < 0.0001, η_p_^2^ = 0.32), and a significant effect of order (F(1,68) = 8.87, p = 0.0040, η_p_^2^ = 0.11), as well as a significant interaction between VNS condition and condition order (F(1,68) = 6.99, p = 0.010, η_p_^2^ = 0.09). Older adults had a small decrease in dynamic range with verum conditions in both condition order groups, while younger adults had an increase in the verum-first group and a decrease in the sham-first group; however, none of the effects of VNS on dynamic range survived post hoc tests.

### 3.7 Effect of VNS on phasic pupillary responses

We fit a linear mixed-effects model on pupillary response maxima relative to baseline (for bright trials, pupillary response minima) with fixed effects of VNS condition, condition order, trial type, age group, and their interactions, and fixed effect of pretrial baseline to account for tonic pupil size drift, with a random slope for VNS condition and random intercept for participant ID to account for the crossover design. The model was evaluated using joint tests.

As expected based based on our analysis in 3.3, there was significant main effect of trial type (F(2,Inf) = 5608.95, p < 0.0001, η_p_^2^ = 0.51)), and an interaction between trial type and age group (F(2,Inf) = 38.79, p < 0.0001, η_p_^2^ = 0.007). We also found a significant maini effect of pretrial baseline such that larger baseline pupil sizes resulted in smaller task-evoked responses (F(1,Inf) = 1113.210, p < 0.0001, η_p_^2^ = 0.12). There was a marginal interaction between VNS condition and trial type (F(1,Inf) = 2.56, p = 0.075, η_p_^2^ = 0.0005), and between trial type, age group and order (F(2,Inf) = 3.35, p = 0.035, η_p_^2^ = 0.0006) and trial type, VNS condition and order (F(2,Inf) = 18.32, p = 0.0001, η_p_^2^ = 0.0017). Following up with pairwise tests between marginal means, we found that verum VNS only significantly affected one trial category: oddball response maxima, while not affecting responses to standard or bright stimuli. This held only in older adults, decreasing phasic response magnitude (z ratio = 2.37, p = 0.018, d = 0.17, see Fig. 4).

**Figure 4.**
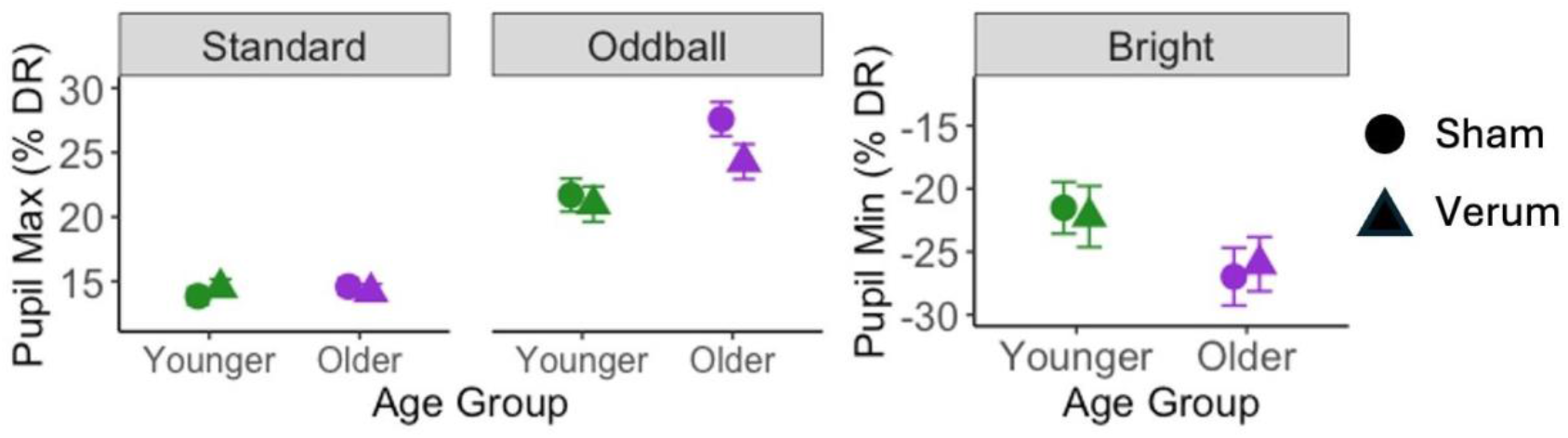
Phasic response maxima (for standard and oddball trial types) and minima (for bright trial type), expressed as percent of individual dynamic range, by age group and VNS condition. Circles show sham stimulation, and triangles show verum stimulation. Error bars show standard error of the mean.

### 3.8 Dose-response effect of current

Since the level of current applied was adjusted to the maximum tolerated by each individual in each condition, we investigated whether the effects of VNS condition on phasic pupillary response depended on the current used. This analysis was limited to oddball trials. In older adults, we fit a linear mixed-effects model on oddball pupillary response maxima with fixed effects of VNS condition and current and their interaction, pretrial baseline pupil size, with a random slope for VNS condition and a random intercept for participant ID. There was a significant interaction between VNS condition and current (F(1,16.23) = 6.19, p = 0.024, η_p_^2^ = 0.30). To determine the effect of current, we plotted estimated oddball response maxima at three levels of current for both sham and verum VNS (Fig. 5). For sham, response maxima did not change with current, but for verum VNS, response maxima decreased as current increased, representing a dose-response effect.

**Figure 5.**
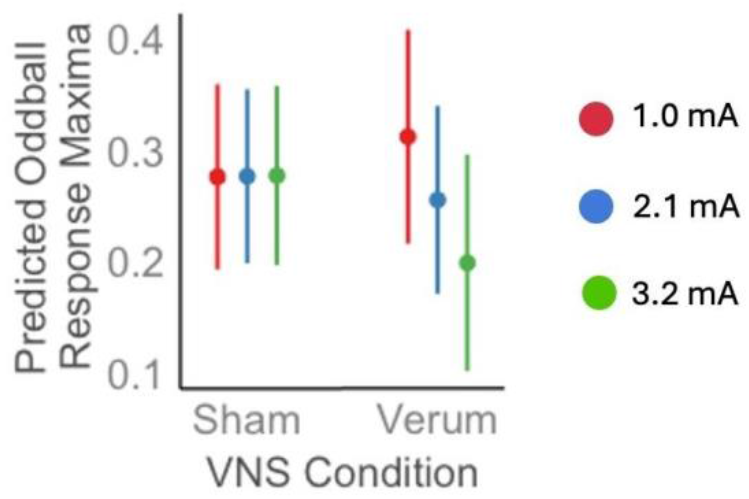
Dose-response relationship between VNS current and pupillary response in older adults. Estimated oddball response maxima (model described in 3.8) at three equally-spaced current levels for sham and verum VNS conditions. Error bars represent 95% confidence intervals.

We repeated this analysis in younger adults. There was also no interaction between VNS condition and current (F(1,15.25) = 0.005, p = 0.94). In a full model with both age groups, there was a significant interaction between age group, VNS condition and current (F(1,33.97) = 5.14, p = 0.048, η_p_^2^ = 0.12), confirming the difference between groups.

Since condition order had a significant effect in other analyses, we also repeated the above models with an effect of condition order and its interactions with VNS condition and current. There was no effect of condition order (F(1,17.21) = 0.09, p = 0.76), nor any significant interactions between condition order and VNS condition or current in older adults (all p > 0.35).

## 4 Discussion

This study is the first investigation of how transcutaneous auricular VNS differentially affects pupillary responses in younger versus older adults. Our findings suggest age-specific responses to VNS that have implications for understanding LC function across the lifespan and for the therapeutic application of VNS to neurodegenerative disease.

Under sham conditions, we replicated established age-related differences in pupillary measures, finding smaller baseline pupil diameter and dynamic range but larger task-evoked responses (expressed as percentage of dynamic range) in older adults compared to younger adults (44). This pattern is consistent with age-related myosis, which may be of central or peripheral origin. These findings align with prior work demonstrating pupillary manifestations of altered LC function in aging and provide further support for their use. We did not, in this study, find evidence for a nonlinear (e.g. quadratic, or inverted-U (55)) relationship between baseline pupil diameter and task-evoked responses. Instead, we only observed a negative linear relationship.

In this study, VNS produced a significant effect in older adults while having minimal impact on younger adults. Specifically, older adults showed increased tonic pupil size and reduced oddball-evoked pupillary responses under verum compared to sham conditions. This effect was subject to a dose-response relationship, such that older adults for whom we used more current had a great reduction in oddball-evoked responses, in the verum condition only. This provides compelling evidence for a physiologically meaningful effect.

VNS responsiveness may differ between age groups because of underlying differences in LC status. For example, cellular responses to early tau pathology, or even simply to a large amount of neuromelanin accumulation, may lead to hyperactivity within the LC (18,25,26). The heightened task-evoked pupillary responses we observed in older adults here and in previous work may be one manifestation of this, and within that context, reduced task-evoked pupillary responses could be seen as beneficial. However, with more severe pathology in the LC comes hypoactivity. Since pretangle tau begins to accumulate in the LC remarkably early (by the second or third decade of life in many cases), but only some people go on to develop LC cell death or Alzheimer’s disease, individuals across nearly the whole adult lifespan could theoretically have LC hyperactivity or hypoactivity. Accurately detecting these states is a very important goal. It is possible that VNS responsiveness may one day serve as a functional diagnostic, allowing researchers or clinicians to infer underlying LC status. Here, younger adults, presumably operating closer to optimal LC activity, appeared to be less responsive to VNS-induced modulation, while in older adults, VNS resulted in tonic pupil size and task-evoked responses that were more similar to those of younger adults.

Several limitations must be acknowledged. The washout period differed between age groups because cognitive testing was conducted during that time and took longer in older adults, although the fact that older adults had both longer washout periods and stronger VNS effects argues against this being a confounding factor. The significant order effects observed in younger adults suggest that future studies should employ a multi-day design to better isolate VNS effects from time-related changes in physiological state. Additionally, our relatively small sample size limits the generalizability of findings, particularly for detecting subtle effects in younger adults.

Most critically, longitudinal studies are needed to determine whether VNS can beneficially impact cognitive aging trajectories. While there are compelling arguments for reducing LC hyperactivity in older adults who may exhibit persistent activation due to cellular damage, there are also theoretical grounds for enhancing LC activity in cases where depression or inattention reflect hypoactivity. In fact, VNS is already FDA approved as a treatment for depression, a condition associated with LC hypoactivity and reduced structural integrity, including in Alzheimer’s disease (56,57). Resolving these competing hypotheses will be essential for developing VNS as a therapeutic intervention for cognitive decline and neurodegenerative disease prevention.

## Acknowledgements

We would like to thank Makanakaishe Chikundura and Zachary Steinbrun for their efforts in data collection. We would also like to extend our appreciation to all the study participants.

